# Long-term respiratory mucosal immune memory to SARS-CoV-2 after infection and vaccination

**DOI:** 10.1101/2023.01.25.525485

**Authors:** Elena Mitsi, Mariana O. Diniz, Jesús Reiné, Andrea M. Collins, Ryan Robinson, Angela Hyder-Wright, Madlen Farrar, Konstantinos Liatsikos, Josh Hamilton, Onyia Onyema, Britta C. Urban, Carla Solorzano, Teresa Lambe, Simon J. Draper, Daniela Weiskopf, Alessandro Sette, Mala K. Maini, Daniela M. Ferreira

**Affiliations:** Oxford Vaccine Group, Department of Paediatrics, University of Oxford, Oxford, UK; Department of Clinical Science, Liverpool School of Tropical Medicine, Liverpool, UK; Division of Infection and Immunity and Institute of Immunity and Transplantation, UCL, London, UK; Department of Biochemistry, University of Oxford, Oxford, UK; Chinese Academy of Medical Science (CAMS) Oxford Institute (COI), University of Oxford, Oxford, UK; Center for Infectious Disease and Vaccine Research, La Jolla Institute for Immunology (LJI), La Jolla, USA; Department of Medicine, Division of Infectious Diseases and Global Public Health, University of California, San Diego, La Jolla, USA

**Keywords:** human lung mucosa, hybrid immunity, SARS-CoV-2 vaccination, airway memory T cells

## Abstract

Respiratory mucosal immunity induced by vaccination is vital for protection from coronavirus infection in animal models. In humans, SARS-CoV-2 immunity has been studied extensively in blood. However, the capacity of peripheral vaccination to generate sustained humoral and cellular immunity in the lung mucosa, and how this is influenced by prior SARS-CoV-2 infection, is unknown. Bronchoalveolar lavage samples obtained from vaccinated donors with or without prior infection revealed enrichment of spike-specific antibodies, class-switched memory B cells and T cells in the lung mucosa compared to the periphery in the setting of hybrid immunity, whereas in the context of vaccination alone, local anti-viral immunity was limited to antibody responses. Spike-specific T cells persisted in the lung mucosa for up to 5 months post-vaccination and multi-specific T cell responses were detected at least up to 11 months post-infection. Thus, durable lung mucosal immunity against SARS-CoV-2 seen after hybrid exposure cannot be achieved by peripheral vaccination alone, supporting the need for vaccines targeting the airways.

## INTRODUCTION

Respiratory mucosal surfaces are the primary site of interaction between SARS-CoV-2 and the immune system. Mucosal antibodies and tissue-resident memory T (TRM) and B cells provide frontline early responses, contributing to protection against the establishment of viral infection following previous viral exposure or vaccination ^1, 2 3^. Animal studies of influenza virus infection have shown that development of antigen-specific resident memory B cells in the lung produces local IgG and IgA with enhanced cross-recognition of variants^4^ and correlates with protection against reinfection in mice^5, 6^. Additionally, studies of respiratory viral infections in animals and human controlled challenge have highlighted the essential role of local tissue-memory T cells in promoting immunity against influenza and respiratory syncytial virus (RSV) mediated, at least in part, by rapid IFN-γ production^7, 8, 9, 10, 11, 12^. Interestingly, in a murine model of SARS-1 and MERS coronavirus infection, protection was attributed to the induction of CD4^+^ T cells in the airway ^13^.

The difficulty accessing human mucosal sites, particularly the lower airways, and the low recovery of fluid and cell yield, have hindered the study of local immunity to respiratory pathogens. Most of the human studies have assessed antibody and T cell responses to SARS-CoV-2 in blood, which is often not reflective of the responses in the airways. However, we and others have demonstrated the presence of pre-existing T cells that recognize SARS-CoV-2 in the lower airways or oropharyngeal lymphoid tissue of unexposed individuals respectively^14, 15^, likely induced by seasonal coronavirus infections. Presence of SARS-CoV-2 specific T cells were also reported in human nasal^16^, lung mucosa and lung-associated lymph nodes following SARS-CoV-2 infection^17,18^. Furthermore, increased numbers of global CD4^+^ and CD8^+^ in the airways of SARS-CoV-2-infected patients were associated with reduced disease severity^19, 20^. It has also been reported that spike-specific memory B cells were enriched in the lungs and associated lymph nodes of convalescent organ donors^18^ and that SARS-CoV-2-binding IgA antibodies are produced more rapidly than IgG and can be detected in the serum and saliva of COVID-19 patients up to 40 days post onset of symptoms ^21, 22, 23, 24^.

Recent animal work with different SARS-CoV-2 vaccine formulations showed the need for mucosal immunisation to generate resident virus-specific B and T cells in the lungs and confirmed the importance of localised mucosal immunity in control of infection ^17, 25, 26, 27^. Human studies that described the effect of peripheral SARS-CoV-2 vaccines on the respiratory mucosa are conflicting. While nasal and salivary IgA responses^28^, as well as CD4^+^ and CD8^+^ T_RM_ were detected in the nasal mucosa^29^ of vaccinated individuals without history of SARS-CoV-2 infection^28^, other studies reported minimal or lack of humoral and T cell responses in nasal and lung mucosa following peripheral vaccination only^17, 30^. However, such responses were detected in convalescent donors or after breakthrough infection^17, 18, 30, 31^.

Further studies are needed to better characterise immune responses in the airways after infection and/or vaccination and dissect out the influence of hybrid immunity, vaccine type, disease severity, and particularly time since vaccination or infection to address persistence of mucosal immunity. Using bronchoalveolar lavage (BAL) samples collected before the onset of the COVID-19 pandemic, we have previously demonstrated that SARS-CoV-2-cross-reactive T cells can reside in human airways^14^. Here, we tested BAL samples and paired blood from vaccinated donors with or without prior SARS-CoV-2 infection and pre-pandemic control samples. We examined the presence of peripheral and mucosal antibodies and virus-specific B and T cell responses. Spike-specific B cells were detected in the airways of individuals exposed to SARS-CoV-2 up to 11 months previously and virus-specific CD4^+^ and CD8^+^ T cells were more abundant in this group compared to vaccinated uninfected individuals. A better understanding of the breath and longevity of adaptive immunity to SARS-CoV2 in the airways will allow us to harness protective mucosal immunity to inform next generation of SARS-CoV-2 vaccines with potential to block infection and population transmission.

## RESULTS

### Characteristics of study groups

To assess humoral and cellular immune responses in the lung mucosa and blood following SARS-CoV-2 vaccination and hybrid immunity, we collected bronchoalveolar lavage (BAL) fluid and paired blood samples from 7 vaccinees with no history or evidence of SARS-CoV-2 infection (naïve, vaccinated group) and 15 vaccinees who had serologically-confirmed asymptomatic infection or experienced symptomatic infection between 2 to 11 months (56 - 333 days) prior to receiving SARS-CoV-2 vaccination. Vaccinated individuals with asymptomatic and symptomatic infection were combined in one ‘hybrid immunity’ group, due to the lack of obvious difference in immune responses to SARS-CoV-2 antigens. All vaccinated individuals received two doses of either mRNA or adenoviral vector vaccine. We also included a pre-pandemic group (n=11) of unexposed, unvaccinated individuals as controls (Figure 1A). Table 1 summarizes the demographic characteristics of the three study groups and the time of sample collection in relation to infection or last vaccination.

**Figure 1.**
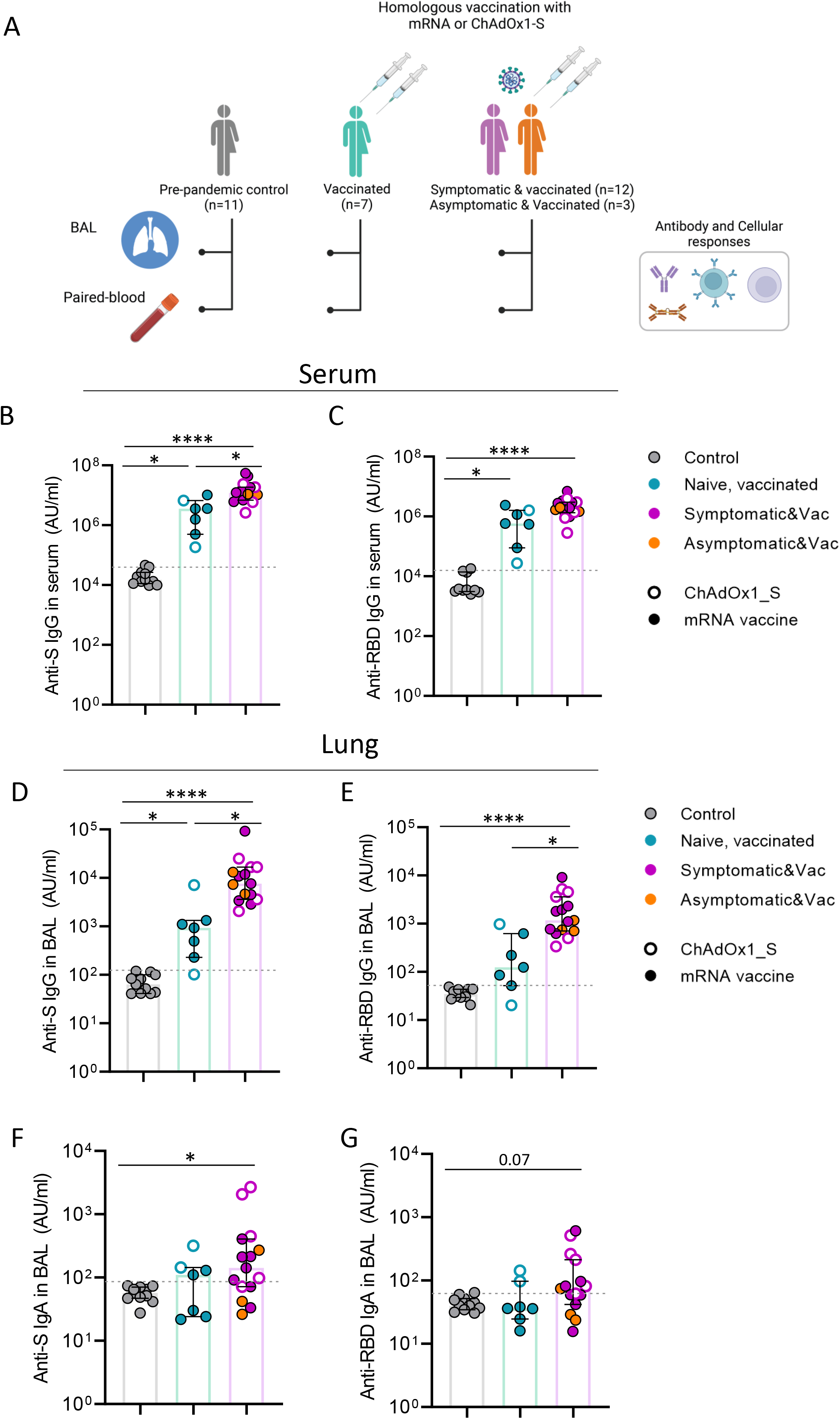
Systemic and lung mucosa antibody responses following vaccination alone and hybrid immunity. **A)** Schematic of study groups depicting SARS-CoV-2 vaccination status, sample collection per group and immunological parameters analysed. Pre-pandemic controls (n=11), infection-naïve vaccinated individual (naïve vaccinated group, n=7) and vaccinated individuals with exposure to SARS-CoV-2 (infected vaccinated or hybrid immunity group, n=15). Different colours used to depict convalescents with asymptomatic or symptomatic SARS-CoV-2 infection. **(B to E)** Levels of IgG against Spike (B and D) and RBD (C and E) in serum and bronchoalveolar lavage (BAL) fluid of control (n=11), vaccinated (n=7) and infected vaccinated donors (n=15). **(F to G)** Levels of IgA against Spike (F) and RDB (G) in BAL fluid of control (n=11), vaccinated (n=7) and infected, vaccinated donors (n=15). Antibody levels are expressed as arbitrary units. Homologous vaccination with ChAdOX1_S or mRNA vaccine is depicted with an open or close circle, respectively. Data shown are in median and interquartile range (IQR). Statistical differences were determined by Kruskal-Wallis test following correction for multiple comparisons. Adjusted p values are indicated by *(p < 0.05), **(p < 0.01) and ****(p < 0.0001).

**Table 1:** Characteristics of participants

| Characteristics | Controls<br>(n=11) | Vaccinated<br>(n=7) | Infected & Vaccinated<br>(n=15) |
| --- | --- | --- | --- |
| Age (yr), median (IQR) | 21 (19 – 24) | 25 (23 – 41) | 43 (20 – 52) |
| Female, n (%) | 7 (64%) | 4 (57%) | 10 (67%) |
| Time since symptomatic<br>infection in days, median (min –<br>max) | n/a | n/a | 273 (201 – 570) |
| NIH clinical score, median (IQR) | n/a | n/a | 4 (3-4) |
| **Vaccination history** |  |  |  |
| Vaccine type | n/a | BNT162b2, n=5<br>ChAdOx1_S, n=2 | BNT162b2, n=9<br>Moderna, n=1<br>ChAdOx1_S, n=5 |
| Time since 2<sup>nd</sup> vaccine dose in<br>days,<br>median (min – max) | n/a | 113 (23 – 186) | 73 (23 -160) |

### Airway antibody responses following hybrid immunity or vaccination alone

Levels of circulating and mucosal antibodies against spike (S), receptor binding domain (RBD) and nucleocapsid (N) protein were measured in serum and BAL samples in all three study groups. Antibody responses to N protein (non-vaccine protein) were used to confirm absence of past infection in the vaccinated cohort and classify the groups. Limit of sensitivity (LOS) was set as median + 2 x standard deviation (SD) of the results in unexposed (pre-pandemic) donors. As expected, anti-N IgG was below or near the LOS in the naïve vaccinated group, whereas in the infected vaccinated group, it was detected in all individuals (15/15) in serum and in 47% (7/15) in the BAL fluid (Extended Data Figure 1A and B).

SARS-CoV-2 vaccination elicited robust systemic IgG responses to both S and RBD protein, with levels being more pronounced in the infected vaccinated group (3.4-fold and 3.3-fold median increase of anti-S and anti-RBD IgG compared to naïve vaccinated group, respectively) (p=0.039) and p=0.11, respectively) (Figure 1B and C). Such systemic antibody differences as a result of hybrid immunity have been extensively demonstrated in large cohort vaccination studies ^32, 33^. High levels of anti-S and -RBD IgG were also detected in the BAL fluid of SARS-CoV-2 vaccinees. Importantly, anti-S and anti-RBD IgG levels in the lung were also significantly elevated in the hybrid immunity group compared to the naïve vaccinated group (8.2-fold and 9.4-fold increase for S and RBD, respectively) (p=0.024 and p=0.014, respectively) (Figure 1D and E).

As IgA plays a crucial role in the antiviral immune defence in mucosal surfaces ^34, 35^, IgA responses against SARS-CoV-2 proteins were also assessed in BAL samples. In the naïve vaccinated group, mucosal IgA levels against S, RBD and N did not differ from the control group. However, the infected vaccinated group had significantly greater mucosal anti-S IgA (2.5-fold increase from control, p=0.014) and a trend to higher anti-RBD IgA (Figure 1F and G), whereas the majority had non-detectable anti-N IgA levels in BAL (Extended Data Figure 1C). Vaccine-induced antibody responses to S protein demonstrated a strong correlation between serum and BAL for IgG and slightly less so for IgA (Extended Data figure 1D and E).

### Presence of SARS-CoV-2 specific memory B cells in the lung following hybrid immunity

Memory B cells are critical for long-term humoral immunity. To identify SARS-CoV-2 specific memory B cells (MBCs), fluorescently labelled S, RBD and N proteins were used to assay PBMCs and lung leukocytes (Figure 2A) (see gating strategy, Extended Data Figure 2A). As expected, and in line with the antibody responses, only vacinees who had previously been exposed to viral nucleoprotein through infection had detectable N-specific MBCs in the blood. By contrast, both naïve vaccinated and infected vaccinated individuals had circulating S- and RBD-specific MBCs above the background staining threshold (set as median + 2 x SD of pre-pandemic levels).

**Figure 2.**
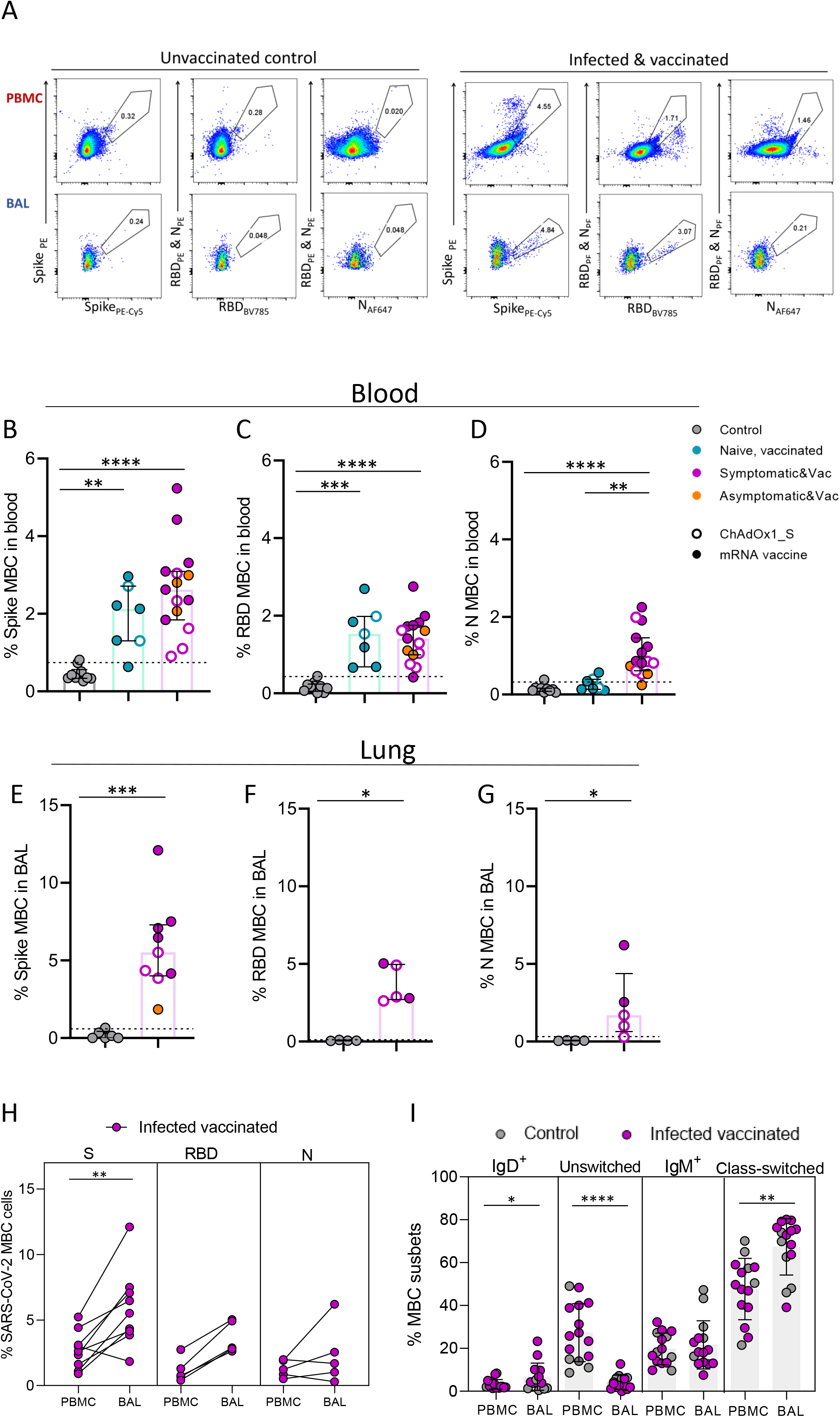
Detectable anti-SARS-CoV-2 memory B cell responses in the lung mucosa following infection and vaccination. **A)** Example flow cytometry plots of Spike-, RBD- and N-specific global memory B cells (MBCs) in PBMC and BAL sample of an unexposed prepandemic control (left) and an infected vaccinated donor(right) (see extended data fig.2 for full gating). **B-D)** Frequency of circulating Spike-, RBD- and N-specific MBCs in control (n=10), naïve, vaccinated (n=7) and infected, vaccinated donors (n=15). E-G) Frequency of Spike-, RBD- and N-specific MBCs detected in BAL samples of control (n=6) and infected vaccinated donors (n=9). H) Frequency of SARS-CoV-2 specific memory B cells in blood (PBMC) and BAL, shown as paired samples, of infected vaccinated donors. (I) Distribution of global MBC subsets in PBMC and BAL based on the expression of IgD and IgM in control and infected, vaccinated donors together (n=25). Homologous vaccination with ChAdOX1_S or mRNA vaccine is depicted with an open or close circle, respectively. Data shown are in median and interquartile range (IQR). Statistical differences were determined by Kruskal-Wallis test following correction for multiple comparisons **(B-D),** Mann-Whitney U test **(E-G)** and Wilcoxon’s paired test **(H-l).** P values are indicated by *(p < 0.05), **(p < 0.01), ***(p < 0.001) and ****(p < 0.0001).

B cells are an underrepresented cell population in the lung mucosa; their presence in the lung is usually associated with infection or chronic inflammation^36^. Although data on anti-viral B cell immunity in human respiratory mucosa are scarce, murine model studies of influenza infection demonstrated the generation of flu-specific memory B cells in the lung following influenza infection that were able to produce more antibodies with enhanced potential to recognise viral variants^4, 5, 6^. In this study, the small number of B cells in the BAL samples allowed the assessment of SARS-CoV-2 specific MBCs only in the hybrid immunity group. The frequencies of S-, RBD- and N-specific MBCs were greater in the lung mucosa of infected vaccinated individuals compared to pre-pandemic controls (median 5.5% vs 0.08% for S, 2.88% vs 0.06% for RBD and 1.69% vs 0.06% for N-specific responses (p=0.0004, p=0.016 and p=0.016, respectively) (Figure 2E–2G). Paired sample comparison of the frequencies of circulating and mucosally detected anti-viral MBCs in the infected vaccinated group revealed enrichment of S- and RBD-specific MBCs in the lung mucosa. The median frequencies of S- and RBD-specific MBCs were 2.1-fold (p=0.0078) and 3.8-fold (p=0.062) higher in the BAL compared to PBMC sample from the same donors (Figure 2H). We detected that in the lung mucosa memory B cells were mainly class-switched MBCs, whereas paired blood samples had a significantly increased proportion of unswitched MBCs cells (Figure 2I).

We also stratified the infected vaccinated group based on the vaccine they received. Despite the small sample size, mRNA vaccinated individuals exhibited 1.8-fold higher frequency of circulating S-specific MBCs compared to ChAdOX1_S recipients (p=0.037), and a trend to higher RBD-specific B cells (Extended Data Figure 2B). In lung mucosa, a similar trend was observed for S-specific MBCs levels (Extended Data Figure 2C), but low cell yields hindered a robust analysis.

### Robust T cells responses in the lung mucosa after infection and vaccination but not vaccination alone

Circulating and tissue resident memory (T_RM_) T cells are important in constraining viral spread and protect against severe disease when neutralising antibodies fail to confer sterilising immunity ^37, 38, 39^. Moreover, we showed that T cells targeting the early expressed replication transcription complex (RTC: NSP7,12,13) are selectively associated with infection being aborted before detection by PCR or seroconversion and can be detected in prepandemic blood and lung samples^14, 40^. Therefore, we examined T cell responses in blood and BAL samples following vaccination alone or infection and vaccination in blood and paired BAL samples.

The frequencies of circulating and lower airway CD4^+^ and CD8^+^ T cells were measured based on the expression of activation-induced markers (AIM assay) after stimulation with SARS-CoV-2 peptides (for full gating strategy see Extended Data figure 3A) and were compared to pre-existing cross-reactive responses detectable using the same assays in cryopreserved pre-pandemic BAL samples. BAL samples were further divided by the expression of prototypic tissue residency markers (CD69/CD103 co-expression for CD8 and CD69/CD49a expression for CD4/CD49a into T_RM_ and recirculating T cells. As reported by others^41, 42, 43^, SARS-CoV-2 vaccination alone induced notable S-specific CD4^+^ and CD8^+^ T cell responses in the circulation when compared to pre-pandemic controls (Figure 3B and 3C). In the infected vaccinated group, the frequency of circulating S-specific CD4^+^ and CD8^+^T cells tended to be higher than the naïve vaccinated group (1.8-fold and 4.8-fold increase, respectively). Despite the induction of T cell immunity systemically, vaccination alone did not elicit S-specific T cell responses that were significantly greater than those in pre-pandemic samples within the global (Figure 3D and 3E) or T_RM_ lung compartment (Figure 3F and 3G).

**Figure 3.**
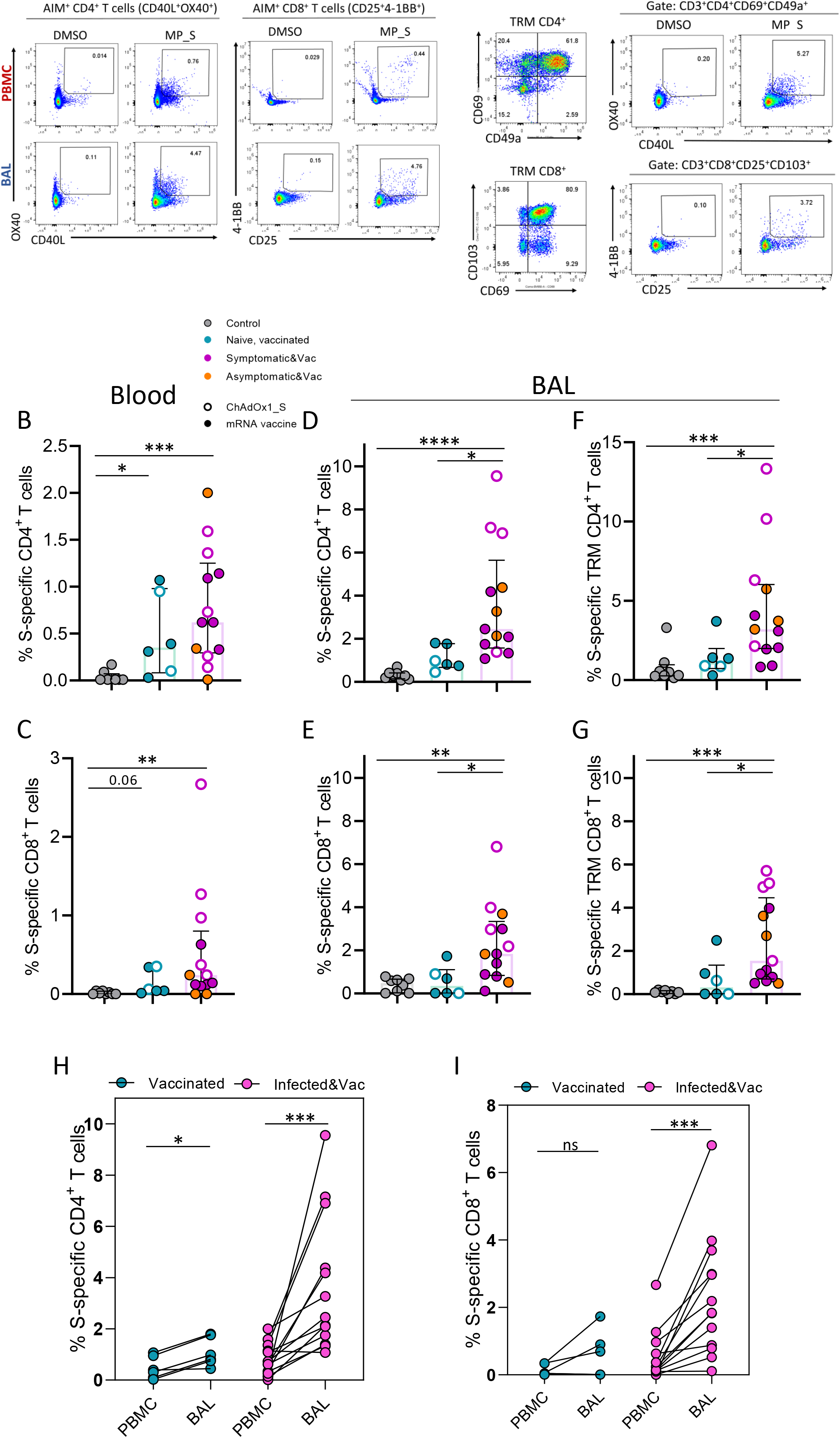
Detection of S-specific T cells responses in the lung mucosa after infection and vaccination but not following vaccination alone. **A)** Representative flow cytometry plots of S-specific CD4^+^ and CD8^+^ T cells in PBMC and BAL sample (on the left) and S-specific TRM CD4^+^ and CD8^+^ T cells in BAL sample (on the right) of an infected, vaccinated donor. Identification of Spike specific T cells was based on the AIM assay, assessing co-expression CD40L and OX40 on CD4^+^ T cells and co-expression of CD25 and 4-1BB on CD8^+^T cells after stimulation with Spike megapools. **B-C)** Frequency of circulating S-specific CD4^+^ and CD8^+^ T cells in control (n=8), naïve, vaccinated (n= 6) and infected, vaccinated donors (n= 13). **D-G)** Frequency of lower-airway S-specific CD4^+^ and CD8^+^ T cells within the global (D-E) and TRM compartment (F-G) in control (n=8), naïve, vaccinated (n= 6) and infected, vaccinated donors (n= 13). H-l) Frequency of S-specific CD4^+^ and CD8^+^ T cells in PBMC and BAL, shown as paired samples, of naïve, vaccinated, and infected vaccinated donors. Homologous vaccination with ChAdOX1_S or mRNA vaccine is depicted with an open or close circle, respectively. Data shown are in median and interquartile range (IQR). Statistical differences were determined by Kruskal-Wallis test following correction for multiple comparisons **(B-G)** and Wilcoxon’s paired test **(H-l).** P values are indicated by *(p < 0.05), **(p < 0.01), ***(p < 0.001) and ****(p < 0.0001).

As opposed to vaccination alone, BAL samples from those who acquired hybrid immunity exhibited greater anti-Spike T cell responses than either the pre-pandemic or naïve vaccinated group (Figure 3D–3G). Within the global T cell population, the frequency of S-specific CD4^+^ and CD8^+^ T cells increased by 2.8-fold and 5.3-fold higher in the infected, vaccinated group compared to the naïve, vaccinated group (p=0.048 and p=0.012, respectively) (Figure 3D and 3E). A similar profile was observed in the T_RM_ T cell compartment, with S-specific CD4^+^ and CD8^+^ T cell frequencies being 2.8-fold and 4.8-fold greater, respectively, in the hybrid immunity group compared to naïve vaccinated group (p=0.05 and p=0.017, respectively) (Figure 3F and 3G). In addition, within the global T cell population, the frequencies of S-specific CD4^+^ and CD8^+^ T cells were substantially higher in BAL than in paired blood from infected vaccinated individuals (median 2.45% vs 0.62% of S-specific CD4^+^ T cells and median 1.84% vs 0.24% of S-specific CD8^+^ T in BAL and paired blood of infected, vaccinated individual, respectively) (p=0.0005 and p=0.0002, respectively) (Figure 3H and 3I). In the naïve, vaccinated group the frequency of S-specific CD4^+^ but not CD8^+^ T cells was slightly higher in BAL than PBMCs (median 0.89% vs 0.35% in BAL and paired blood, respectively).

In agreement with large vaccination studies^44, 45^, we observed that adenoviral vector vaccine tended to induce increased frequency of S-specific T cells in the periphery compared to mRNA vaccines. The tendency of the adenoviral vector vaccine to induce stronger T cell immunity was also observed in the lower airways, with 3.3-fold and 2.5-fold higher S-specific CD4^+^ and CD8^+^ T cell responses when compared to RNA vaccination (Extended Data Figure 3B-C).

We also examined T cell specificities for non-vaccine included SARS-CoV-2 structural proteins (N and membrane [M]) and non-structural proteins (NSP-7, NSP-12 and NSP-13 pool, representative of the core replication-transcription complex [RTC]) in blood and BAL (Extended Data Figure 3D-E and Figure 4). As expected, the frequencies of circulating N- and M-specific CD4^+^ and CD8^+^ T cells were significantly higher than in pre-pandemics only in the hybrid immunity group, as those vaccinees had a past SARS-CoV-2 infection (Figure 4A and 4B). In the case of RTC-specific T cells, their frequency did not differ amongst groups, as SARS-CoV-2 cross-reactive CD4^+^ and CD8^+^ T cell responses were detected systemically in 3 out 8 pre-pandemic controls, in line with previous studies^14, 40, 46^. In BAL samples, the frequency of the aforementioned T cell specificities was tested in a subset of pre-pandemic and infected, vaccinated individuals based on cell number availability. Interestingly, the hybrid immunity group had, or tended to have, higher N- and M- and RCT-specific T cell responses within the global and TRM T cell compartment in BAL samples compared to levels detected in pre-pandemic controls (Figure 4C–4F). In addition, these SARS-CoV-2 specific CD4^+^ and CD8^+^ T cell responses were enriched in the lower airways compared to the periphery (Figure 4G and 4H).

**Figure 4.**
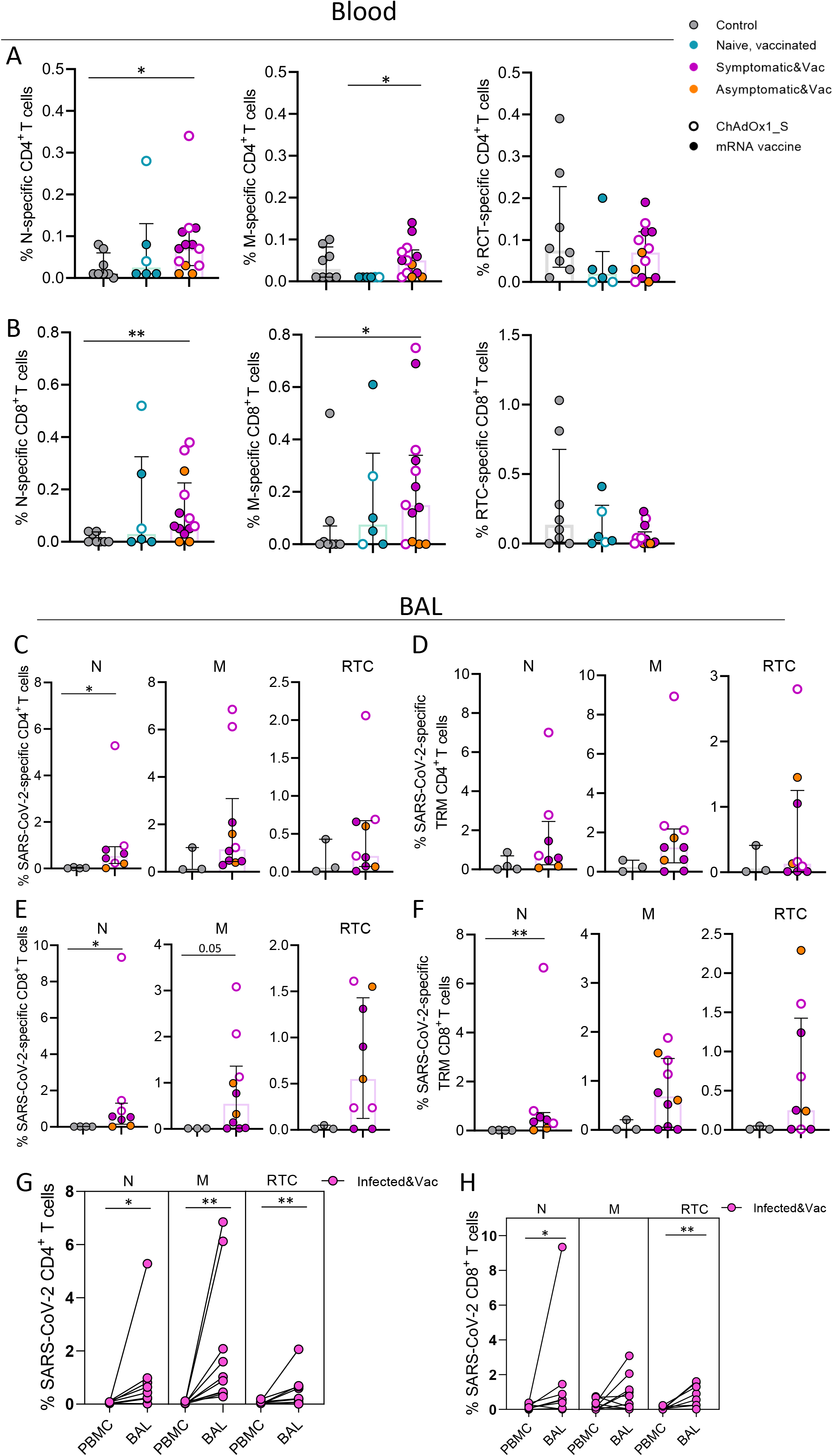
Detection of infection-induced SARS-CoV-2 T cell responses in the periphery and lung mucosa. **A-B)** Frequency of circulating N-, M- and RTC-specific CD4^+^ and CD8^+^ T cells in control (n=8), naïve, vaccinated (n= 6) and infected, vaccinated donors (n= 13). **C-F)** Frequency of N-, M- and RTC-specific CD4^+^ and CD8^+^ T cells within the global and TRM compartment of lower airway T cells in control (n=4) and infected, vaccinated donors (n=8). **G-H)** Frequency of SARS-CoV-2-specific CD4^+^and CD8^+^ T cells in PBMC and BAL, shown as paired samples, of infected, vaccinated donors. Homologous vaccination with ChAdOX1_S or mRNA vaccine is depicted with an open or close circle, respectively. Data shown are in median and interquartile range (IQR). Statistical differences were determined by Kruskal-Wallis test following correction for multiple comparisons **(A-B),** Mann-Whitney U test **(E-F)** and Wilcoxon’s paired test (G-H). P values are indicated by *(p < 0.05) and **(p < 0.01).

The hierarchy of SARS-CoV-2 antigen recognition by circulating and lower airway T cells of each distinct peptide pool (S, N, M and RTC) was analysed in a subset of 8 infected vaccinated individuals (Extended Data Figure 3D-E). The antigen recognition profile differed between systemic and airway localised T cells, and between T cell subsets. SARS-CoV-2 specific CD4^+^ T cells were largely dominated by S-specific CD4^+^T cells in the periphery and lung mucosa, however in lower airway they were enriched with additional T cell specificities (Extended Data figure 3D). In the case of SARS-CoV-2 CD8^+^ T cells, their antigen recognition profile was more diverse in both sites, with S-specific CD8^+^ T cells being apparent but not dominant (Extended Data figure 3E).

### Longevity of antibody and T cells responses in lung mucosa following vaccination alone or hybrid immunity

To assess the longevity of vaccine-induced SARS-CoV-2 immune memory in the lung mucosa following vaccination and hybrid immunity, antibody and T cell responses assessed in BAL and paired blood of naïve, vaccinated and infected vaccinated individuals were plotted in association with time post the 2^nd^ vaccine dose (which was 2-11 months after any known infection dates). Levels of circulating anti-S and anti-RBD IgG were negatively correlated with time post-vaccination in the naïve vaccinated but not the infected vaccinated group, implying quicker antibody decay in the naïve vaccinated donors (Figure 5A). In the lung mucosa, anti-S and anti-RBD IgG levels exhibited similar rates of decay between the two vaccinated groups (Figure 5B). On the other hand, levels of anti-S and RBD IgA in BAL were detectable only following hybrid immunity but they quickly reached pre-pandemic levels (at 5-months post-vaccination, Figure 4C). This result is in agreement with previous studies in convalescent patients that reported short-lived IgA-mediated immunity at mucosal sites^21, 47^.

**Figure 5.**
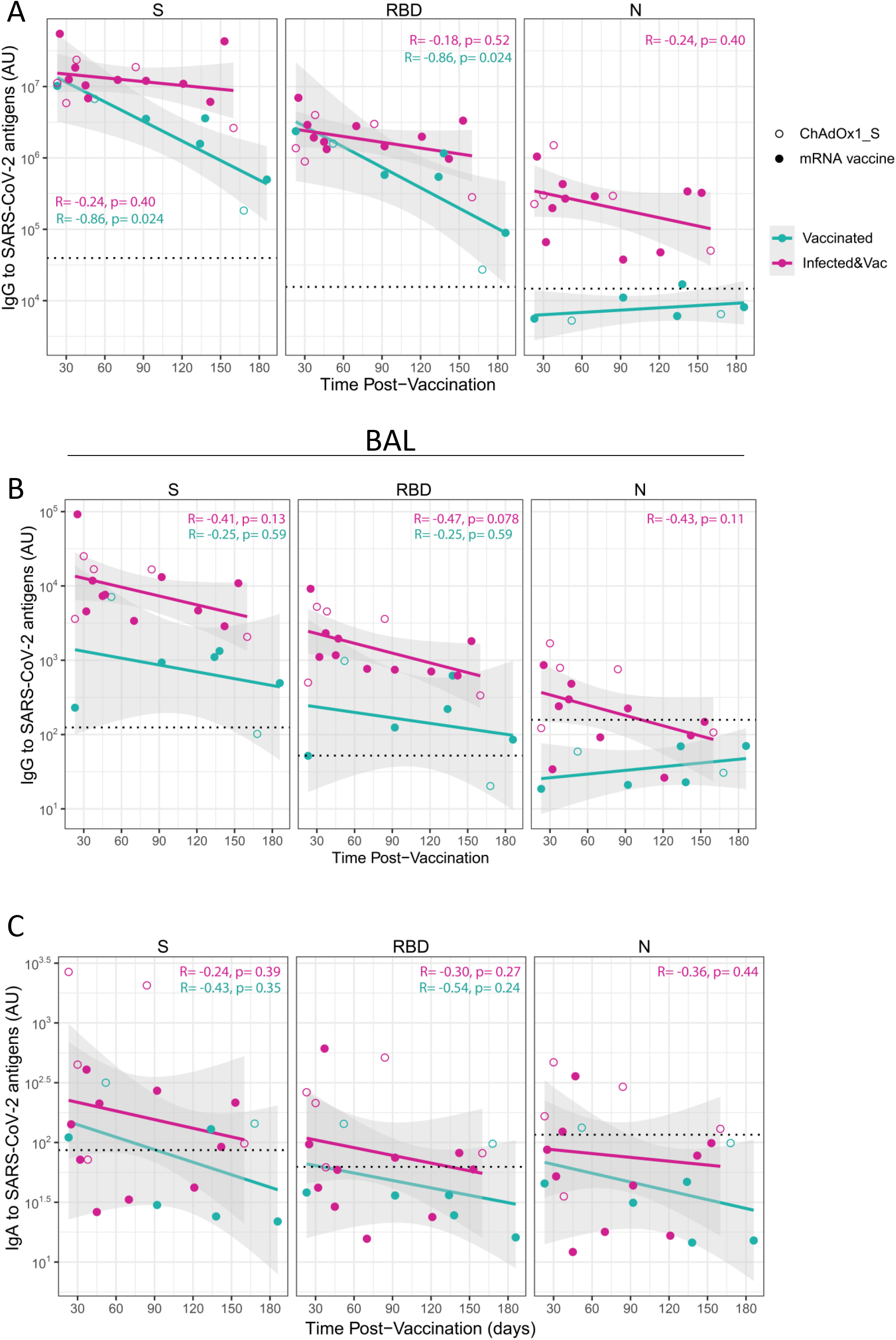
Long-lived IgG but short-term IgA responses to SARS-CoV-2 antigens in the lung mucosa following vaccination and infection. **A-B)** Correlation between time postvaccination and levels of IgG to S, RBD and N proteins measured in serum (A) and BAL supernatant (B) of naïve vaccinated (n=7) and infected vaccinated individuals (n=15). **C)** Correlation between time post-vaccination and levels of IgA to S, RBD and N proteins measured in BAL supernatant of of naïve, vaccinated (n=7) and infected, vaccinated individuals (n=l5). Homologous vaccination with ChAdOX1_S or mRNA vaccine is depicted with an open or close circle, respectively. Spearman correlation used. R and p values are shown.

Circulating and lower-airway S-specific T cell frequencies were also plotted in association with time post-vaccination in both vaccinated groups. In blood, numbers of S-specific CD4^+^ and CD8^+^ T cells declined over time in the infected vaccinated group and at 5-months post-vaccination reached the frequencies induced by vaccination alone (Figure 6A). On the other hand, S-specific T cell responses were better sustained in the lung mucosa following hybrid immunity. Despite a trend of negative association with time post-vaccination, S-specific CD4^+^ T cell responses were detectable from the lung mucosa of infected vaccinated individuals for over 5-months post vaccination. Lower-airway S-specific CD8^+^T cells did not associate negatively with time, suggesting they remained at stable levels throughout the period of 5-months post vaccination (Figure 6B and 6C). The human lung also retained partial immune memory to SARS-CoV-2 over a year post infection. Despite decline over time, T cell specificities not affected by SARS-CoV-2 vaccination, such as M-, N- and RTC-specific CD4^+^ and CD8^+^ T cells, were detectable in various frequencies between 6 to 18 months post infection (Figure 6D), rendering these conserved SARS-CoV-2 proteins as potential vaccine targets.

**Figure 6.**
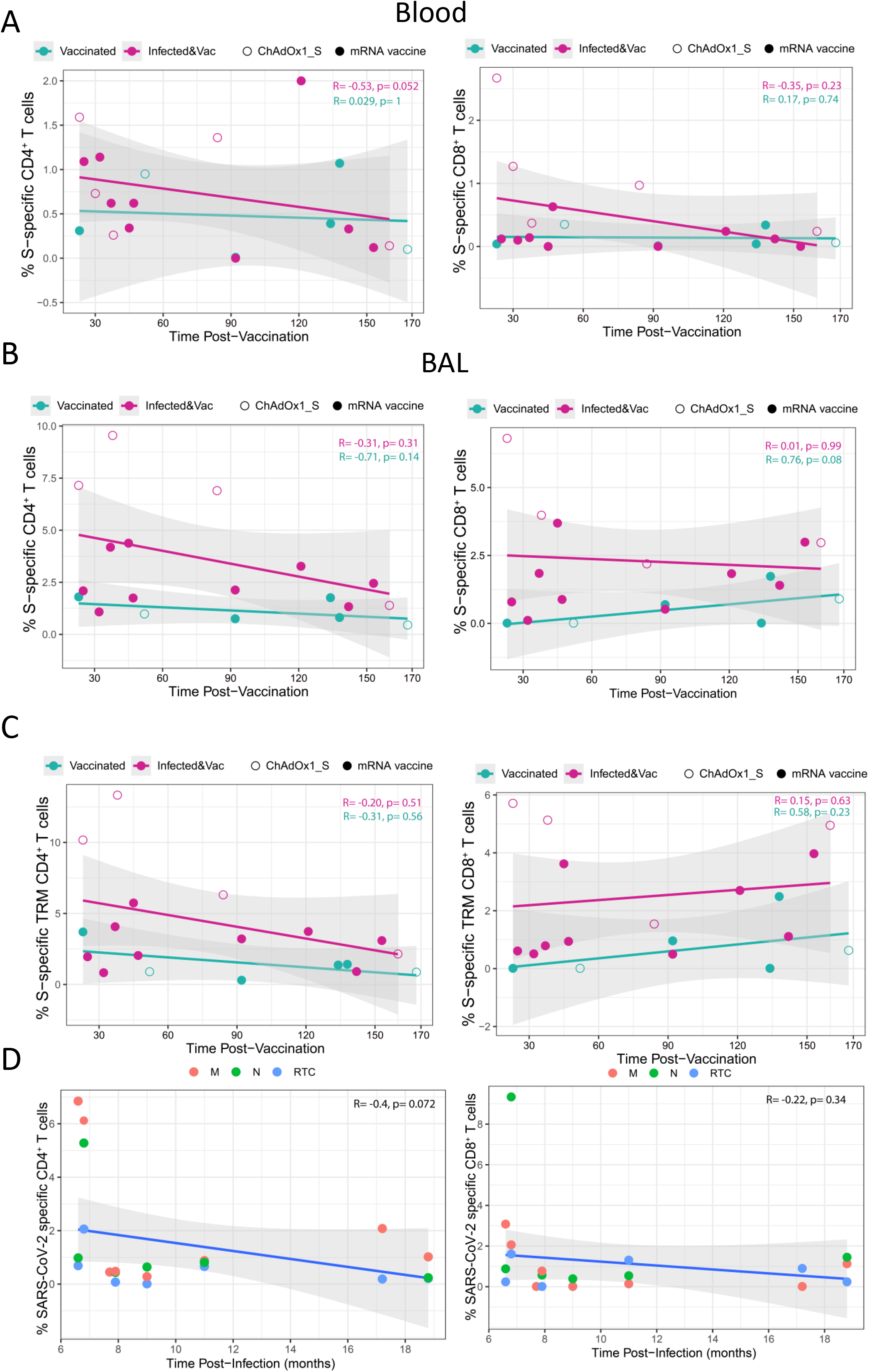
Persistent S-specific T cells in the lung mucosa following hybrid immunity. **A)** Correlation between time post-vaccination and the frequency of S-specific CD4^+^ (left) and CD8^+^ T cells (right) detected in the blood of naïve vaccinated (n=6) and infected vaccinated individuals (n=15). **B-C)** Correlation between time post-vaccination and the frequency of S-specific CD4^+^ (left) and CD8^+^ T cells (right) in the global and TRM compartment of lower-airway T cells in BAL of naïve, vaccinated (n=6) and infected, vaccinated individuals (n=15). Homologous vaccination with ChAdOX1_S or mRNA vaccine is depicted with an open or close circle, respectively. D) Correlation between time post infection from symptoms onset and the frequency of N-, M and RTC-specific CD4^+^ (left) and CD8^+^ T cells (right) detected in BAL of infected, vaccinated individuals with symptomatic, PCR-confirmed SARS-CoV-2 infection (n=13). Spearman correlation used. R and p values are shown.

## DISCUSSION

Despite high SARS-CoV-2 seroprevalence globally by either vaccination or infection, regular waves continue to cause breakthrough infections. Immunological memory to SARS-CoV-2 in the respiratory mucosa following vaccination, infection and hybrid immunity is not well understood. Protective mucosal immunity could be harnessed for the development of vaccines specifically targeting protection against airway infection to block transmission of the virus in the population.

Here, we assessed the potential of peripheral SARS-CoV-2 vaccination to induce anti-viral immune memory in the human lung mucosa and whether a prior infection would influence the immunological outcome. We found that following SARS-CoV-2 vaccination alone the lung mucosa was enriched with spike IgG, but levels were increased and accompanied by mucosal IgA in the setting of hybrid immunity. Importantly, homologous parenteral SARS-CoV-2 vaccination (mRNA or adenoviral vector vaccine) did not seed the human lung with tissue-residing spike-specific T cells, despite the induction of notable T cell responses in the circulation. Compared to SARS-CoV-2 vaccination alone, hybrid immunity resulted in considerably higher humoral and cellular immune responses against the vaccine antigens in the periphery. In contrast to vaccination alone, hybrid immunity generated robust and persistent spike-specific T cell immunity in the human lung mucosa, complemented with local MBC and T cell responses against additional SARS-CoV-2 antigens. A long-lived, airway-compartmentalised B and T cell reservoir in the lung mucosa may confer better recognition of Omicron sublineages and future variants and protect against severe disease, supporting the need for vaccines specifically targeting the airways.

In line with others^17, 31^, our results indicate that following vaccination alone, SARS-CoV-2 immunity in the respiratory mucosa is limited to humoral immunity, with IgG dominating over IgA titres against the vaccine-antigens. Induction of both anti-Spike IgG and IgA was more efficient in the lung mucosa of vaccinated individuals with prior infection. The strong correlation observed between anti-S IgG in serum and BAL samples supports the notion that systemic antibodies elicited by vaccination transudate to the respiratory mucosa, as previous vaccination studies have reported^48, 49^. Despite the key role of antibodies in neutralising the virus at the respiratory mucosa-the primary site of infection, local humoral immunity wanes quickly^21^ making individuals more susceptible to immune escape by Omicron sublineages and future variants^17^. In addition, findings from other respiratory infection and vaccination studies indicate that higher levels of antibodies are required in the nasal mucosa to protect against local infection compared to levels required in blood to protect against invasive disease ^50^. However, the finding of class-switched memory B cells enriched in the lung mucosa raises the possibility they produce a repertoire of antibodies better able to cross-recognise variants, as shown in other infections^4^. Mucosal antibodies may also harness local lung cells such as NK cells and phagocytes for non-neutralising Fc-dependent cellular immunity.

Booster parenteral vaccination is required to enhance waning humoral immunity, but the frequency and intensity of robust systemic T-cell responses is not boosted by the additional vaccination^51^. Hybrid immunity elicits considerably high humoral and cellular responses in the periphery, which exceed the immune responses induced solely by vaccination^32, 52, 53^. Studying the human lung mucosa, we provide the first evidence, to our knowledge, that hybrid immunity, contrary to SARS-CoV-2 immunisation alone, can generate robust, broader, and long-lived anti-viral immune responses in the lower airways. In line with other studies^17, 18^, the intensity of S-specific B and T cell responses were enriched in the lung mucosa compared to the periphery and similarly, enriched B and T cells responses were detected for other SARS-CoV-2 antigens. These lower-airway localised B and T cells reservoirs were long-lived. Hence using more comprehensive analysis of T cell specificities, we demonstrated that lower airway T cell immunity against SARS-CoV-2 can be sustained for over a year post the initial exposure to the pathogen. In studies of other respiratory viruses, lung localised, tissue-residing B and T cells associate with protection in mouse models of influenza^5,54^ and RSV infection^10^. In human challenge models of influenza and RSV infection, enrichment of CD4^+^ and CD8^+^ T_RM_ cells in the airways was associated with mitigated respiratory symptoms, viral control, and reduced disease severity^12, 55^. The prolonged memory together with the ability of T cells to better recognise more conserved parts of SARS-CoV-2, support the utility of developing multi-specific mucosally-administered vaccines that could boost tissue localised and resident memory T and B cells in the lung. Preclinical studies of SARS-CoV-2 have demonstrated that intranasal vaccination decreases viral shedding and transmission relative to parenteral vaccines^27, 56, 57^. In addition, the combined approach of systemic priming SARS-CoV-2 vaccination followed by intranasal boosting with either adenovirus vectored vaccines or an adjuvanted Spike vaccine elicited both systemic and protective mucosal immunity with cross-reactive properties^58, 59^. In humans, there are limited data on the immunogenicity of SARS-CoV-2 vaccines that target the airways, focusing mainly on humoral immunity^60^.

Our study has limitations. Due to the invasive nature of the bronchoscopy procedure, we were not able to recruit a large cohort of study participants. In addition, the fast roll out and uptake of C0VID-19 vaccine programme in the UK hindered the recruitment of convalescent unvaccinated individuals. The low BAL cell yields restricted the analysis of other T cell specificities to selected SARS-CoV-2 proteins and did not allow the assessment of vaccine-induced memory B cells in the lung mucosa of infection naïve individuals. Given the inability of SARS-CoV-2 parenteral immunisation to generate tissue-localised T cell immunity, it is less likely that S-specific memory B cells will be present in the lung mucosa following vaccination. Although, we were able to detect and characterise T cell specificities for over a year post infection and up to 6 months post vaccination, future longitudinal studies are needed to fully understand the long-term impact of airway localised T and B cell immunity in SARS-CoV-2 protection.

Overall, our data suggest robust lung mucosal immunity against SARS-CoV-2 can be better achieved through hybrid immunity, as opposed to peripheral vaccination alone. Vaccines that induce airway localised memory T and B cells may provide broader long-term protection at the site of infection. Thus, vaccination approaches that combine systemic and mucosal immunisation could reduce virus transmission and re-infection cases.

## METHODS

### Study design and cohorts

This was a cross-sectional study, which included a cohort of SARS-CoV-2 vaccinated individuals (n=22), who had received two doses of mRNA or the ChAdOx1_S adenoviral vector vaccine. A subset of them had experienced PCR-confirmed symptomatic infection (n=12) or serologically confirmed asymptomatic infection (n=3), referred to the group of infected and vaccinated individuals (hybrid immunity) (n=15), whereas the remaining vaccinees had not experienced SARS-CoV-2 infection (vaccinated, n=7). BAL samples were obtained through research bronchoscopy 1 to 6 months (23-186 days) after the 2^nd^ vaccine dose and 7 to 19 months (201 -570 days) after symptoms onset for those who had experienced COVID-19. These COVID-19 convalescent individuals had been admitted to hospital between April 2020 to January 2021, when the ancestral SARS-CoV-2 strain was still dominant in the UK. Blood samples for sera and PBMC isolation were collected at the same day as BAL. Pre-pandemic samples from healthy, unexposed individuals (n=11), collected from 2015 to 2018, were also included into the analysis, as a control group (Figure 1A). The demographic and clinical characteristics of the 3 study groups are shown in Table 1.

### Sample processing

BAL samples were processed as previously described^61^, cryopreserved in CTL-CryoABC medium kit (Immunospot). After thawing, alveolar macrophages were routinely separated from other non-adherent immune cell populations using an adherence step, as previously described^62^. Blood was processed for sera collection or PBMCs were isolated from heparinized blood samples using density-gradient sedimentation layered over Ficoll-Paque in SepMate tube and then cryopreserved in CTL-CryoABC medium kit (Immunospot).

### ELISA for SARS-CoV-2 proteins

ELISA was used to quantify levels of IgG and IgA against Spike trimer, RBD and N in serum and BAL samples, as previously described^63^. Briefly, 96-well plates (U bottom) were coated with 1μg/ml SARS-CoV-2 antigen and stored at 4°C overnight for at least 16h. The next day, plates were washed 3 times with PBS/0.05% Tween-20 and blocked with 2% BSA in PBS for 1h at room temperature. Sera and BAL diluted in 0.1% BSA-PBS were plated in duplicate and incubated for 2h at room temperature alongside an internal positive control (dilution of a convalescent serum) to measure plate to plate variation. For the standard curve, a pooled sera of SARS-CoV-2 infected participants was used in a two-fold serial dilution to produce either eight or nine standard points (depending on the antigen) that were assigned as arbitrary units. Goat anti-human IgG (γ-chain specific, A9544, Millipore-Sigma) or IgA (α-chain specific, A9669, Millipore-Sigma) conjugated to alkaline phosphatase was used as secondary antibody, and plates were developed by adding 4-nitrophenyl phosphate in diethanolamine substrate buffer. Optical densities were measured using an Omega microplate reader at 405nm. Blank corrected samples and standard values were plotted using the 4-Parameter logistic model (Gen5 v3.09, BioTek).

### B cells immunophenotyping and detection of SARS-CoV-2 specific B cells

Cryopreserved BAL cells and PBMCs were used for detection of SARS-CoV-2 specific B cells in lower airways and blood, respectively. Biotinylated tetrameric S, RBD and N protein were individually labelled with different streptavidin conjugates at 4°C for 1h, as previously described^63^.

Biotinylated S and RBD were directly labelled with Streptavidin-PE (with a ratio 1:3 and 1: 5.7, respectively); with Streptavidin-BV570 (S with a ratio 1: 2.7); and Streptavidin-BV785 (RBD with a ratio 1:5). Biotinylated N protein was labelled with Streptavidin-PE (with a ratio of 1:2.3) and Streptavidin-AF647 (N protein with a ratio 1: 0.5).

PBMCs and BAL cells were thawed and stained with Live/dead e506 viability dye and an antibody cocktail for surface markers for 30min in the dark, washed twice and resuspended in 200μL of PBS. Parallel samples stained with an identical panel of monoclonal antibodies (mAbs),but excluding the SARS-CoV-2 proteins (fluorescence minus one [FMO] controls), were used as controls for nonspecific binding. All samples were acquired on an Aurora cytometer (Cytek Biosciences) and analysed with Flowjo software version 10 (Treestar). The flow-cytometry panel of mAbs used to phenotype global and antigen-specific subsets can be seen in Supplementary Table 1.

The frequency of antigen-specific B cells was calculated within the fraction of MBCs (CD19^+^CD27^+^, excluding the naïve IgD^+^CD27^-^ and the double negative IgG^-^CD27^-^ fractions, see gating strategy (Extended Data Figure 2A). For phenotypic analysis of spike-, RBD-, and N-specific B cells, a sufficient magnitude of responses (≥50 cells in the relevant parent gate) was required.

### Activation-induced markers (AIM) T cell assay

Mononuclear BAL cells (1 × 10^5^ cells per well) and PBMCs (1 × 10^6^ cells per well) were seeded in 96-well plates in RPMI supplemented with 1% PNS and 10% AB human serum (Merck, UK) and stimulated with SARS-CoV-2 specific peptides pools. The peptides pools used were spanning the whole Spike protein based on predicted epitopes (15-mer peptides overlapping by 10 amino acids)^64^ or overlapping peptides spanning the immunogenic domains of the SARS-CoV-2 N (Prot_N) and M (Prot_M) purchased from Miltenyi Biotec^63^ or combined pools spanning SARS-CoV-2 NSP7, NSP12 and NSP13 proteins (15-mer peptides overlapping by 10 amino acids) of the ancestral SARS-CoV-2 strain^14,65^. Prior to the peptide addition, cells were blocked with 0.5μg/ml of anti-CD40 mAb (Miltenyi Biotec) for 15min at 37°C. A stimulation with an equimolar amount of DMSO was performed as a negative control and Staphylococcal enterotoxin B (SEB, 2 μg/mL) was included as a positive control. The following day cells were harvested from plates, washed and stained for surface markers (Supplemental table 2 and 3).

AIM^+^CD4^+^ T cells were identified as CD40L^+^OX40^+^, 4-1BB^+^OX40^+^, 4-1BB^+^CD40L^+^ subsets, and the CD40L^+^OX40^+^ combination was used to quantify SARS-CoV-2 specific CD4^+^ T cells frequency. SARS-CoV-2 specific CD8^+^ T cells were identified as 4-1BB^+^CD25^+^. Antigen-specific CD4^+^ and CD8^+^ T cells were measured and presented as DMSO background-subtracted data.

### Statistical analysis

Participant characteristics were summarised as n, median (interquartile range) or frequency (percentage). Chi-squared test and Fisher’s exact test were conducted to identify any significant changes in categorical variables. Non-parametric Wilcoxon paired tests and Mann-Whitney tests were conducted to compare quantitative data within the same group or between two groups, respectively. In addition, Kruskal-Wallis rank sum test with Dunn’s correction were performed to compare quantitative data amongst groups (three groups comparison). All tests were two-sided with an α level of 0.05. To explore the association between time after infection and vaccination, we employed a linear regression model. Data were analysed in R software version 4.0.3 (R Foundation for Statistical Computing, Vienna, Austria) or Graphpad Prism version 9.0.

## Ethics statement

All volunteers gave written informed consent and research was conducted in compliance with all relevant ethical regulations. Ethical approval was given by the Northwest National Health Service Research Ethics Committee (REC no. 18/NW/0481 and Human Tissue licensing no. 12548).

## Acknowledgements

We thank all the patients and healthy volunteers who participated in the present study and all the clinical staff who helped with recruitment and sample collection. The study was financially supported by EPSRC grant (no. EP/W016389/1) awarded to E.M and D.M.F and a Centre of Excellence in Infectious Diseases Research (CEIDR) pump-priming grant. Collection of pre-pandemic clinical samples was supported by the Bill and Melinda Gates Foundation (grant no. OPP1117728) and the UK Medical Research Council (grant no. M011569/1) awarded to D.M.F. The production of peptide pools has been funded or in part with federal funds from the National Institutes of Health, Contract No. 75N9301900065 to A.S, D.W.

M.K.M. is supported by Wellcome Trust Investigator Award (no. 214191/Z/18/Z) and CRUK Immunology grant (no.26603).

## Authors contribution

E.M, M.O.D, M.K.M and D.M.F conceived and designed the study. A.H.W and M.F recruited and consented study participants. A.H.W, M.F, R.R and A.M.C obtained human samples. E.M, J.R, J.H and CS processed samples. E.M, M.O.D, J.R, J.H, O.O and T.L generated and analysed the data. E.M. M.O.D, M.K.M and D.M.F interpreted data. E.M, J.R and B.U developed the assays. S.J.D, D.W and A.S provided material for the assays. E.M and M.O.D prepared the manuscript. All authors provided critical input into the manuscript.

---

**Extended Data Figure 1.**
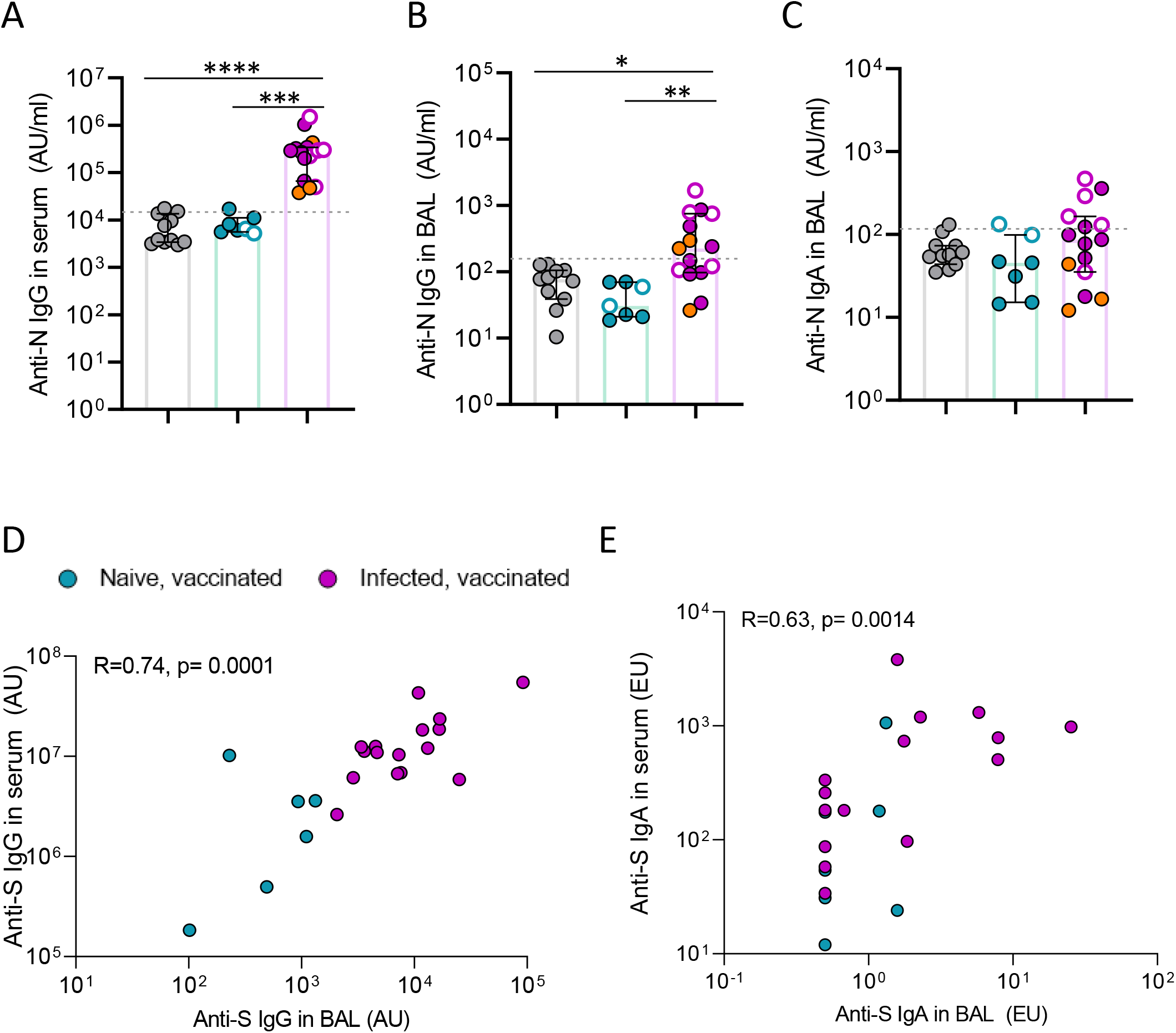
Associations between lung mucosa and systemic antibody levels against Spike post SARS-CoV-2 vaccination. **(A to C)** Levels of anti-N IgG in serum (A) and anti-N IgG (B) and IgA (C) in BAL fluid of control (n=11), naïve vaccinated (n=7) and infected vaccinated donors (n=15). Correlation of anti-S IgG (D) and anti-S IgA (E) levels measured in serum and BAL fluid of naïve, vaccinated (in turquoise) and infected, vaccinated donors (in purple). Total n=22 vaccinated individuals. R and p values are shown using Spearman rank correlation.

**Extended Data Figure 2.**
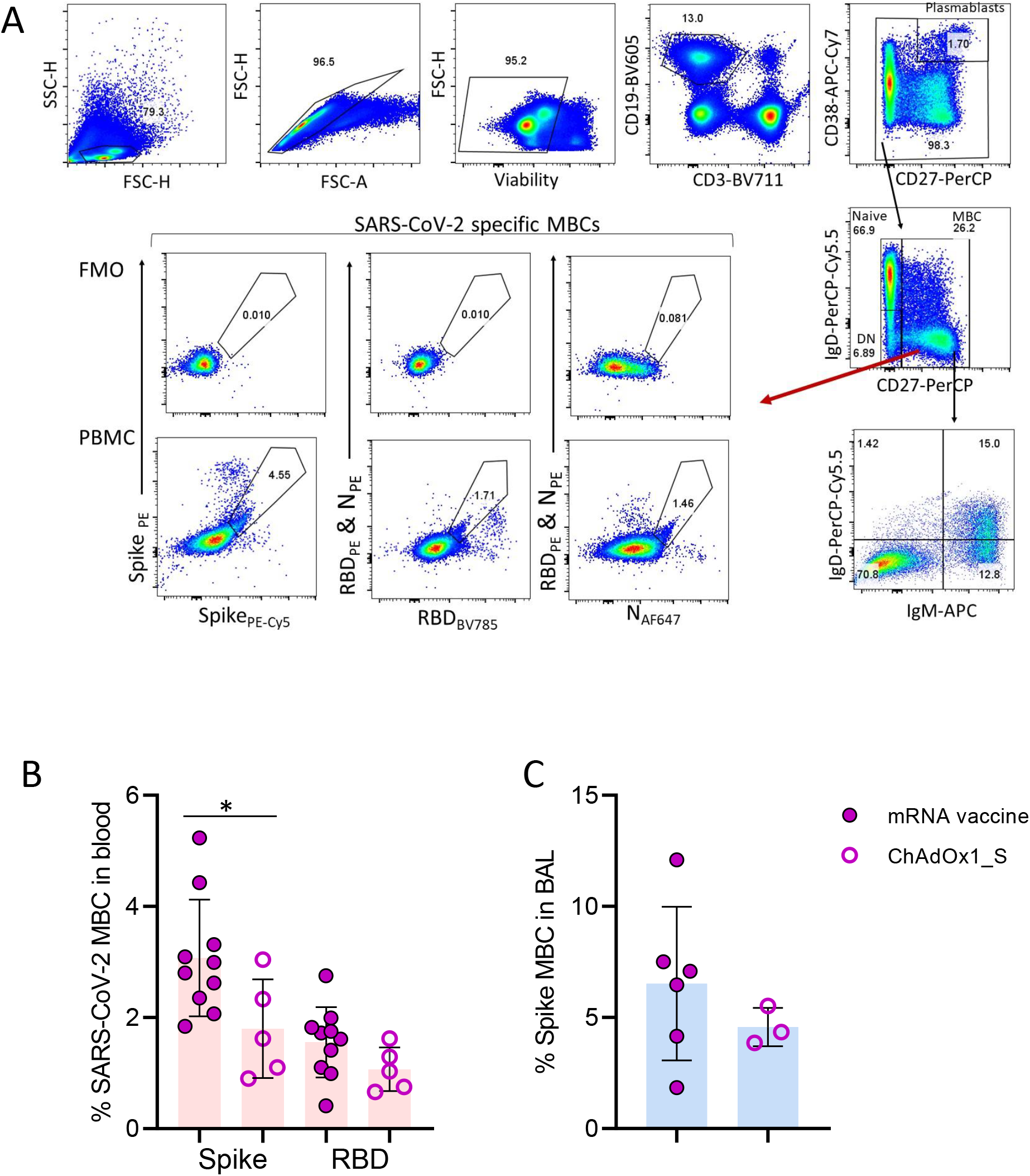
Memory B cells and SARS-CoV-2 specific B cells responses in human blood and BAL. **A)** Gating strategy of B cells in human PBMC or BAL sample, with representative flow cytometry plots of SARS-CoV-2 specific B cells from controls and infected, vaccinated donors. B) Frequency of circulating SARS-CoV-2 MBCs in mRNA (n=10) or ChAdOxAd1_S vaccine (n=5) recipients with prior infection. C) Frequency of lower airway Spike MBCs in mRNA (close circles, n=6) or ChAdOxAd1_S vaccine (open circles, n=3) recipients with prior infection. Data shown are in median and interquartile range (IQR). Statistical differences were determined by Mann-Whitney U test. P values are indicated by *(p < 0.05), **(p < 0.01) and ****(p < 0.0001).

**Extended Data Figure 3.**
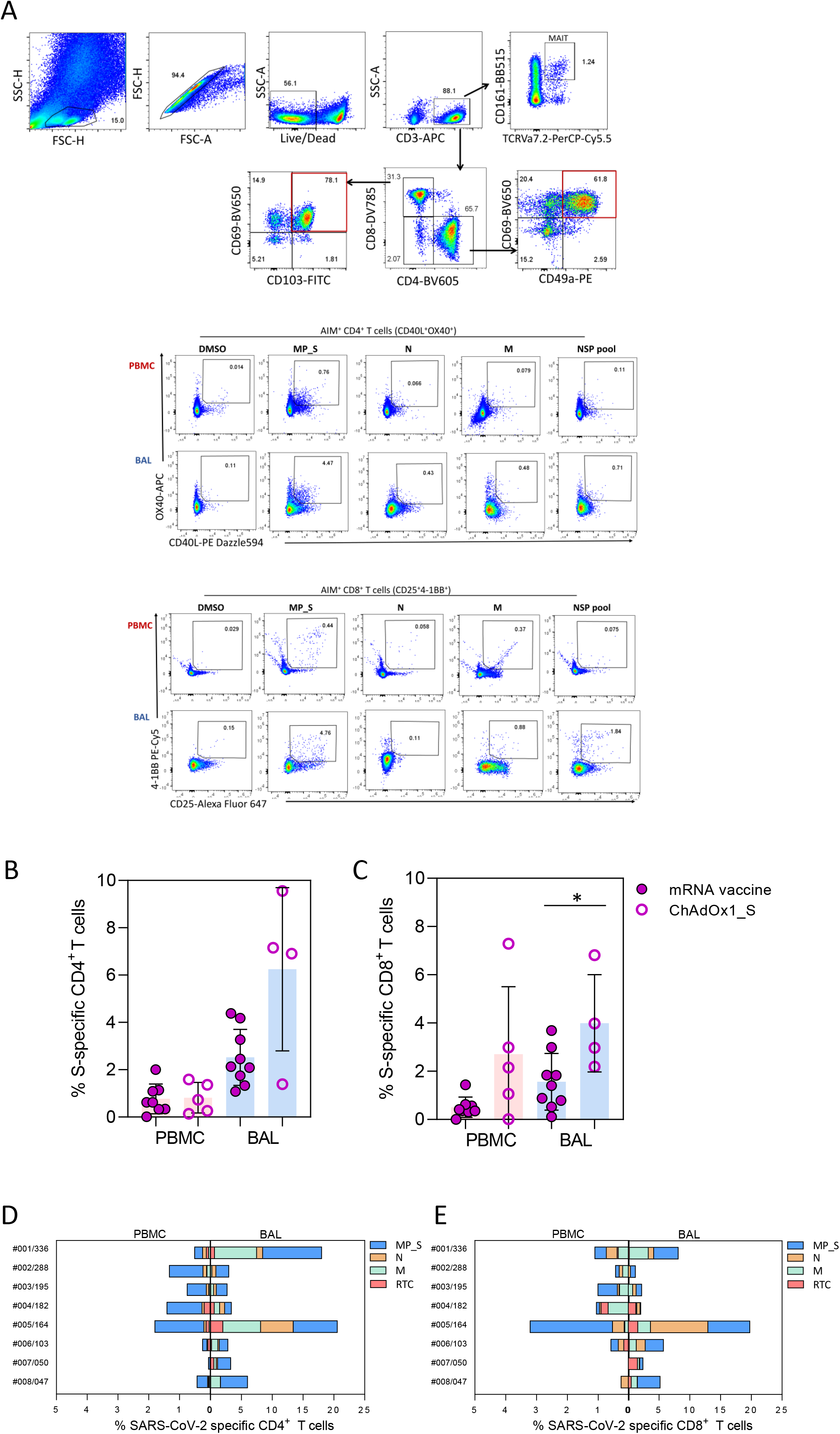
**A)** Example gating of T cell staining after overnight stimulation with SARS-CoV-2 peptide pools. Lymphocytes (SSC-H vs. FSC-H), single cells (FSC-H vs. FSC-A), Live cells (fixable live/dead), CD3^+^, CD4^+^ or CD8^+^ or MAIT cells. Tissue resident memory (TRM) CD4^+^ and CD8^+^ T cells were defined as CD4^+^CD69^+^CD49a^+^ and CD8^+^CD69^+^CD103^+^, respectively. B) Representative plots of SARS-CoV-2 specific CD4^+^ and CD8^+^ T cells in PBMC and BAL of an infected, vaccinated donor after overnight stimulation with SARS-CoV-2 peptide pools. DMSO was used as a negative control. **B-C)** Frequency of circulating and lower-airway S-specific CD4^+^ and CD8^+^ T cells in mRNA (n=9) or ChAdOxAd1_S vaccine (n=5) recipients with prior infection. **D-E)** Paired analysis of proportions of CD4^+^ and CD8^+^ T cells which recognise SARS-CoV-2 antigens in BAL and paired blood. Statistical differences were determined by Mann-Whitney U test. *(p < 0.05).

