## Supplementary material for "Long-term respiratory mucosal immune memory to SARS-CoV-2 after infection and vaccination"

| **Marker** | **Fluorochrome** | **Clone** | **Provider** | **Reference** |
| --- | --- | --- | --- | --- |
| CD21 | SB436 | HB5 | Thermofisher | 62-0219-42 |
| CD95 (Fas) | BV421 | DX2 | Biolegend | 305624 |
| CD11c | Pacific Blue | Bu15 | Biolegend | 337212 |
| CD185 (CXCR5) | BV480 | RF8B2 | BD Biosciences | 566142 |
| CD19 | BV605 | HIB19 | Biolegend | 302244 |
| CD20 | BV650 | 2H7 | Biolegend | 302336 |
| CD3 | BV711 | SK7 | Biolegend | 344838 |
| CD27 | PE-eFluor610 | O323 | Thermofisher | 61-0279-42 |
| CD24 | FITC | ML5 | Biolegend | 311104 |
| IgD | PerCPCy5.5 | IA6-2 | Biolegend | 348208 |
| CCR10 | PECy7 | 13E11 | Biolegend | 362108 |
| IgM | APC | SA-DA4 | ThermoFisher | 17-9998-42 |
| CD71 | AF700 | M-A712 | BD Biosciences | 563769 |
| CD38 | APC-Cy7 | HIT2 | Biolegend | 303534 |
| Live & Dead | e506 | NA | Thermofisher | 65-0866-14 |

**Supplemental table 1. Summary of antibody cocktail used for B cell immunophenotyping.** A multiparametric flow cytometry panel consisted of 14 monoclonal antibodies and a viability dye was used to immunophenotype global and memory B cells in blood and BAL samples from the 3 study groups. Electronic compensation was set using CompBeads (BD Biosciences), Arc beads (ThermoFisher) and using the Spectral Flow automated unmixing software (Aurora, Cytek) according to manufacturer’s instructions.

| **Marker** | **Fluorochrome** | **Clone** | **Provider** | **Reference** |
| --- | --- | --- | --- | --- |
| CD45RA | BV750 | H100 | BD Biosciences | 747435 |
| CD197 (CCR7) | BV480 | 3D12 | BD Biosciences | 566099 |
| CD3 | APC-Cy7 | SK7 | Biolegend | 344818 |
| CD4 | BV605 | SK3 | Biolegend | 344646 |
| CD8 | BV785 | SK1 | Biolegend | 344740 |
| CD185 (CXCR5) | SB436 | MU5UBEE | Thermofisher | 62-9185-42 |
| CD183 (CXCR3) | BV510 | G025H7 | Biolegend | 353726 |
| CD196 (CCR6) | PECy7 | G034E3 | Biolegend | 353418 |
| TCRVa7.2 | PerCPCy5.5 | 3C10 | Biolegend | 351710 |
| CD161 | BB515 | REA631 | Miltenyi Biotech | 130-122-808 |
| CD69 | BV650 | FN50 | Biolegend | 310934 |
| CD154 (CD40L) | PE/Dazzle594 | 24-31 | Biolegend | 310840 |
| CD137 (4-1BB) | PECy5 | 4B4-1 | Biolegend | 309808 |
| CD134 (OX40) | APC | ACT35 | Biolegend | 350008 |
| CD25 | AF647 | BC96 | Biolegend | 302617 |
| Live & Dead | e506 | NA | Thermofisher | 65-0866-14 |

**Supplemental table 2.** Summary of antibody cocktail used for T cell immunophenotyping and identification of antigen-specific T cells in blood using AIM assay**.**

| **Marker** | **Fluorochrome** | **Clone** | **Provider** | **Reference** |
| --- | --- | --- | --- | --- |
| CD45 | BV750 | HI30 | BD Biosciences | 747321 |
| CD3 | APC-Cy7 | SK7 | Biolegend | 344818 |
| CD4 | BV605 | SK3 | Biolegend | 344646 |
| CD8 | BV785 | SK1 | Biolegend | 344740 |
| CD183 (CXCR3) | BV510 | G025H7 | Biolegend | 353726 |
| TCRVa7.2 | PerCPCy5.5 | 3C10 | Biolegend | 351710 |
| CD161 | BB515 | REA631 | Miltenyi Biotech | 130-122-808 |
| CD69 | BV650 | FN50 | Biolegend | 310934 |
| CD103 | FITC | Ber-ACT8 | Biolegend | 350204 |
| CD49a | PE | TS2/7 | Biolegend | 328304 |
| CD154 (CD40L) | PE/Dazzle594 | 24-31 | Biolegend | 310840 |
| CD137 (4-1BB) | PECy5 | 4B4-1 | Biolegend | 309808 |
| CD134 (OX40) | APC | ACT35 | Biolegend | 350008 |
| CD25 | AF647 | BC96 | Biolegend | 302617 |
| Live & Dead | e506 | NA | Thermofisher | 65-0866-14 |

**Supplemental table 3.** Summary of antibody cocktail used for T cell immunophenotyping and identification of antigen-specific T cells in BAL using AIM assay**.**
